# Myogenic artifacts masquerade as neuroplasticity in the auditory frequency-following response (FFR)

**DOI:** 10.1101/2023.10.27.564446

**Authors:** Gavin M. Bidelman, Alexandria Sisson, Rose Rizzi, Jessica MacLean, Kaitlin Baer

## Abstract

The frequency-following response (FFR) is an evoked potential that provides a neural index of complex sound encoding in the brain. FFRs have been widely used to characterize speech and music processing, experience-dependent neuroplasticity (e.g., learning, musicianship), and biomarkers for hearing and language-based disorders that distort receptive communication abilities. It is widely assumed FFRs stem from a mixture of phase-locked neurogenic activity from brainstem and cortical structures along the hearing neuraxis. Here, we challenge this prevailing view by demonstrating upwards of ∼50% of the FFR can originate from a non-neural source: contamination from the postauricular muscle (PAM) vestigial startle reflex. We measured PAM, transient auditory brainstem responses (ABRs), and sustained frequency-following response (FFR) potentials reflecting myogenic (PAM) and neurogenic (ABR/FFR) responses in young, normal-hearing listeners with varying degrees of musical training. We first establish PAM artifact is present in all ears, varies with electrode proximity to the muscle, and can be experimentally manipulated by directing listeners’ eye gaze toward the ear of sound stimulation. We then show this muscular noise easily confounds auditory FFRs, spuriously amplifying responses by 3-4x fold with tandem PAM contraction and even explaining putative FFR enhancements observed in highly skilled musicians. Our findings expose a new and unrecognized myogenic source to the FFR that drives its large inter-subject variability and cast doubt on whether changes in the response typically attributed to neuroplasticity/pathology are solely of brain origin.

## Introduction

Neuroelectric brain recordings have been indispensable in demonstrating neuroplasticity at all levels of the auditory system. In particular, the frequency-following response (FFR), a scalp-recorded neurophonic reflecting phase-locked activity along the auditory neuroaxis to periodic signals, has served as a neural index of sound coding in the EEG. The precision of the FFR is evidenced by the fact these brain potentials can support intelligible speech when they are replayed (i.e., sonified) as audio stimuli (Bidelman, 2018a). The strength to which FFRs capture voice pitch (i.e., fundamental frequency; F0) and harmonic timbre cues of complex signals is also related to listeners’ perception of speech material (Weiss and Bidelman, 2015). FFRs are also enhanced by various experiential factors including native language experience (Krishnan et al., 2010; Zhao and Kuhl, 2018), musical abilities (Wong et al., 2007; Mankel and Bidelman, 2018), and perceptual learning (Reetzke et al., 2018). Conversely, FFRs reveal deficiencies in neural processing in a variety of auditory, literacy, and neurodevelopmental disorders (e.g., dyslexia, autism, hearing loss, aging; Russo et al., 2008; Chandrasekaran et al., 2009; Anderson et al., 2013; White-Schwoch et al., 2015; Bidelman et al., 2017a). Collectively, a wealth of studies implies FFRs might be a valuable auditory biomarker for tracking both positive and maladaptive auditory plasticity in the brain that either bolsters or compromises the perceptual organization of speech and musical sounds.

Despite an abundance of FFR studies and potential applications to understanding brain plasticity and normal and disordered central auditory processing, the anatomical origins of the FFR remain highly contentious (Coffey et al., 2016; Holmes and Herrmann, 2017; Bidelman, 2018b; Coffey et al., 2019; White-Schwoch et al., 2019). Historically described as a brainstem potential (Smith et al., 1975; Sohmer and Pratt, 1977; Chandrasekaran and Kraus, 2010), it is now recognized FFRs reflect a mixture of phase-locked activity from brainstem and cortical structures throughout the auditory pathway (Bidelman, 2018b; Coffey et al., 2019). While different neuroimaging techniques emphasize brainstem- and cortico-centric contributions to the response, the FFR has always been described, unequivocally, as a *brain response* of *auditory-neurogenic* origin (Hoormann et al., 1992; Chandrasekaran and Kraus, 2010; Bidelman and Powers, 2018). However, anecdotal observations as earlier as the late 1970s (Sohmer et al., 1977)^1^, and our own experience over the past decade reveal occasional listeners that produce unusually large FFRs that far exceed the amplitudes expected for an auditory-neurogenic potential in humans. Such enigmatic responses are easily obscured in grand average data but can differ from the FFRs’ normal operating range (Chandrasekaran and Kraus, 2010) by an order of magnitude (i.e., 1-2 µV vs. 100-200 nV range), and thus warrant further explanation. Here, we expose a heretofore unrecognized *myogenic* source of the FFR that explains substantial individual differences in the response including waveform enhancements usually attributed to plasticity in central auditory nervous system function (Krishnan et al., 2005; Musacchia et al., 2007; Wong et al., 2007; Kraus et al., 2009; Parbery-Clark et al., 2009; Krizman et al., 2012; Kraus et al., 2014; Coffey et al., 2017; Mankel and Bidelman, 2018; Zhao and Kuhl, 2018).

Situated behind the ear, the postauricular muscle (PAM) is part of a vestigial startle reflex that once acted to retract the pinna to protect hearing (Bérzin and Fortinguerra, 1993). The muscle produces large (>100 µV), bilateral contraction with peak latency of 9-15 ms following brief sounds (Thornton, 1975; O’Beirne and Patuzzi, 1999) and is highly variable across listeners (Picton et al., 1974). Anesthetic blockage of the muscle and facial nerve abolishes PAM confirming its non-neural origin (Bickford et al., 1964). While PAM activation declines with increasing stimulus rates (Geisler et al., 1958), the reflex can be elicited by surprisingly fast periodic stimuli without fatigue or habituation (Jacobson et al., 1964) (but see Yoshie and Okudaira, 1969). Indeed, motor units underlying the PAM reflex can be driven at rates up to 100-200 Hz (Kiang et al., 1963; Jacobson et al., 1964), resulting in steady-state muscle potentials that, like the FFR, phase-lock to periodic signals (Kiang et al., 1963). Problematically, these rates closely coincide with the low-F0 (∼100 Hz) pitched stimuli used in most auditory FFR studies (Coffey et al., 2016). This raises the possibility that FFRs to low-frequency sounds, including the voice pitch and low harmonics of speech, might be partially driven by muscular rather than auditory neurophonic structures. This is particularly relevant in the context of typical FFR recording approaches, which place electrodes behind the ear where PAM reflex is optimally recorded (O’Beirne and Patuzzi, 1999). Indeed, in our informal survey of FFR studies published in the past 50 years (N=314 studies), nearly half (42%=131 papers; see *SI Materials*) used a mastoid reference electrode montage that promotes extraneous pickup of PAM muscle artifact (**Fig. 1a**). The problem thus appears widespread in the literature. More critically, identifying undocumented muscular confound in the FFR would be particularly germane to interpreting plasticity studies and the unchallenged assumption that amplified FFRs (e.g., as in musicians, bilinguals) necessarily reflect increased fidelity of auditory neural coding due to listening experience (cf. Musacchia et al., 2007; Wong et al., 2007; Kraus et al., 2009; Parbery-Clark et al., 2009; Bidelman et al., 2011; Krizman et al., 2012; Coffey et al., 2017; Mankel and Bidelman, 2018; Zhao and Kuhl, 2018).

**Figure 1:**
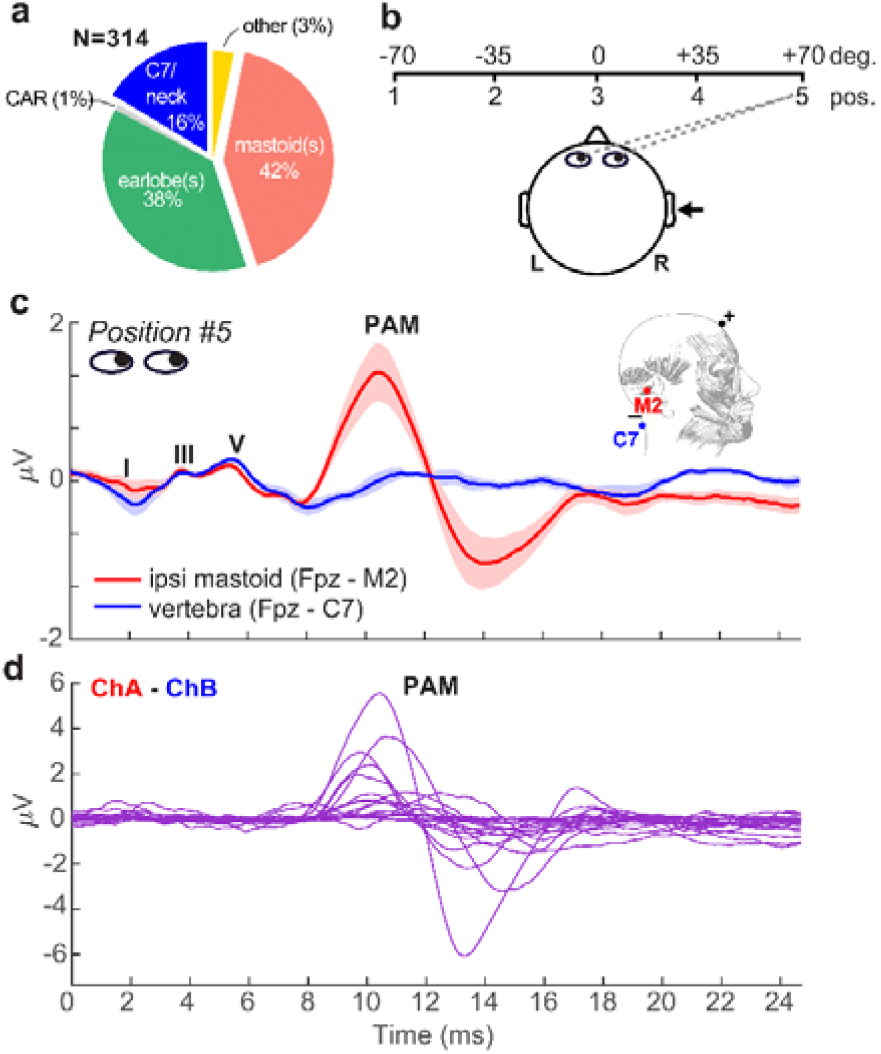
Eye gaze toward the ear introduces PAM artifact in ABRs. (**a**) In our literature review of N=314 FFR studies, nearly half (42%=131 papers) use a mastoid reference electrode montage that can promote inadvertent pickup of muscle artifact (see *SI Materials*). (**b**) Schematic of the visual gaze paradigm to direct listeners’ eyes during ABR/FFR recording. Listeners were positioned 1 m from the booth wall and were cued to direct their eye gaze to one of 5 positions spanning ±70 degrees from head center relative to the ipsilateral ear of stimulus presentation (right ear). **(c)** Grand average ABR waveforms recorded with a vertical ipsilateral montage between a non-inverting electrode at the high forehead (∼Fpz) referenced to an inverting electrode placed on the (i) ipsilateral mastoid (M2) or (ii) 7^th^ cervical vertebra (C7). Eye gaze was directed to position 5 (70 deg) in both cases. Strong PAM is recorded for the channel with electrode over the ipsilateral mastoid. Moving the reference to C7 (i.e., distal to the PAM muscle) eradicates pickup of the artifact. (**d**) Individual data, with PAM responses isolated via subtraction of the traces in panel C. PAM strength ranges from 0.5-10 _µ_V (peak-to-peak) across individuals. Anatomy adapted from Feneis (1994). Shading = ± 1 s.e.m.

Among the more widely reported neuroplastic effects captured in the FFR are the putative enhancements reported in musicians (Schneider et al., 2002; Musacchia et al., 2007; Wong et al., 2007; Kraus and Chandrasekaran, 2010). Speech-FFRs are stronger and shorter in latency in musicians relative to their nonmusicians peers (Kraus et al., 2009; Parbery-Clark et al., 2009), providing a neural account of their enhanced speech perception observed behaviorally. Such enhancements are typically interpreted as reflecting stronger neural representation for speech shaped by experience-dependent plasticity from the enriched sonic environment afforded by musical engagement (Wong et al., 2007; Kraus and Chandrasekaran, 2010; Herholz and Zatorre, 2012). However, recent studies have challenged this notion by demonstrating even nonmusicians with superior music-listening abilities (i.e., “musical sleepers”) have “enhanced” FFRs that mirror those of highly trained musicians (Mankel and Bidelman, 2018). Such findings reveal innate differences in auditory brain function (and possibly other unmeasured factors) can easily masquerade as plasticity in studies on the brain benefits of music (Musacchia et al., 2007; Wong et al., 2007; Tierney et al., 2015). The present study was not intended to refute the possible connections between musicianship and FFR enhancements observed in both cross-sectional and longitudinal studies (Musacchia et al., 2007; Wong et al., 2007; Tierney et al., 2015; Nan et al., 2018). Rather, we aimed to identify whether PAM muscle activity naturally induced by sound stimulation might not only be larger in musicians but partially mediate the auditory processing enhancements reported in studies on music-related plasticity.

To this end, we evaluated whether unmeasured muscle artifact might explain substantial inter-subject variability in the FFR and account for at least some of the neural enhancements in speech processing frequently reported in highly skilled listeners (e.g., musicians). A major source of variation in the strength of PAM elicitation stems from uncontrolled eye movements (Patuzzi and O’Beirne, 1999). Gaze directed toward the ear of sound presentation potentiates PAM contraction and thus pickup of the muscle artifact in EEG (O’Beirne and Patuzzi, 1999; Patuzzi and O’Beirne, 1999). By experimentally manipulating PAM with a gaze paradigm, we show here that repeated stimulation of the muscle produces “following-like” potentials in the EEG via simple linear superposition of overlapping PAM wavelets (cf. Bidelman, 2015). Moreover, we find this muscle noise easily masquerades as the auditory FFR and spuriously amplifies the response by 3-4x fold, partially accounting for FFR enhancements observed in highly skilled musicians. As a solution for recording artifact-free FFRs, we further show PAM-FFR contamination is easily circumvented by using high-frequency stimuli and relocating electrodes to a non-cephalic (neck) site.

## Materials and Methods

### Participants

The sample included *N* = 20 young adults (age [*µ ± σ*]: 24.9 ± 2.8 years; 4 male, 16 female). This sample size was determined *a priori* to match comparable studies on auditory plasticity and the FFR (Parbery-Clark et al., 2009; Bidelman et al., 2014; Coffey et al., 2016). All had normal hearing (i.e., pure-tone air-conduction thresholds ≤ 25 dB HL; 250-8000 Hz), similar levels of education (19.1 ± 2.3 years), and reported no previous history of neuropsychiatric illness. All but one individual were right-handed (77 ± 45% laterality) (Oldfield, 1971). The sample included a range of formal musical training (*µ ± σ*: 6.5 ± 7.3 years; range 0-23 years) to assess whether putative enhancements in the FFR reported in experienced listeners (cf. Musacchia et al., 2007; Wong et al., 2007; Skoe and Kraus, 2012; Mankel and Bidelman, 2018) might instead result from undocumented PAM muscle artifact rather than auditory neuroplasticity, *per se*. Music training was treated as a continuous rather than binary variable since there is disagreement on what constitutes the definition of a “musician” (Zhang et al., 2020) and we used self-report (as in previous FFR studies) rather than a formal tests of music listening skills (cf. Mankel and Bidelman, 2018). Each gave written informed consent in compliance with a protocol approved by Indiana University IRB.

### Stimuli

Auditory brainstem responses (ABRs) were recorded to 100 µs clicks. Frequency-following responses (FFRs) were elicited by complex pitch stimuli (periodic click trains) with fundamental frequencies (F0s) of either 100 or 200 Hz (Bidelman, 2015). Pulse trains were constructed using a periodic series of impulses 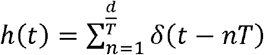, where *d* is the duration of the desired pulse train (here, 100 ms), and *T* is the period between successive impulses (i.e., 1/F0). Each pulse of the train was constructed using identical clicks (bandwidth, pulse with, amplitude) to those described for eliciting the ABR. This allowed us to directly test the assumption that a large amplitude periodic FFR response is a linear superposition of repeated transient PAM waves (cf. Bidelman, 2015).

### EEG recordings

Participants sat in an electro-acoustically shielded booth in a recliner chair positioned 1 m from the wall. Numbered signs (labeled 1-5) were positioned in the front hemifield demarcating angles of -70, -35, 0, +35, +70 degrees (left to right) relative to head center (see **Fig. 1b**)^2^. For each condition, we directed participants’ eye gaze to one of the five angles by cueing the respective number. Compliance was monitored by the experimenter via a window into the chamber. FFRs were recorded with eye gaze directed at each of the 5 azimuths.

FFRs were recorded using a two-channel vertical montage with Ag/AgCl disc electrodes placed on the mid-hairline (i.e., midway between Fpz and Fz) referenced to (i) right (ipsi) mastoid (M2) and (ii) 7^th^ cervical vertebra (mid-forehead= ground). Due to technical error, data from the second channel were not recorded in N=2 listeners and were treated as missing values. Impedances were ≤ 5 kΩ. EEGs were digitized at 20 kHz using a SmartEP EEG system (Intelligent Hearing Systems; Miami, FL) with an online passband of 50 -3000 Hz (+ 60 Hz notch filter) and 100K amplifier gain. Evoked responses were elicited from each participant in response to right ear presentation at 80 dB SPL through electromagnetically shielded insert earphones (Ultra-shielded ER3A inserts, 300Ω; IHS, Miami, FL) that eradicated electromagnetic stimulus artifact from contaminating biological responses (Price and Bidelman, 2021). Stimuli were presented with fixed, rarefaction polarity at a repetition rate of 9.09/s. Presentation order was randomized both within and across participants. Continuous EEGs were epoched (*ABR*: 0-24.7 ms; *FFR:* 0-127 ms) and ensemble averaged across trials to derive evoked responses per condition. Sweeps > ±50 µV were automatically rejected during online averaging. A total of 2000 artifact-free sweeps were collected per stimulus and eye gaze position.

### Model simulating FFRs from PAM artifact

To evaluate the degree to which FFRs are explained by PAM artifact, we compared the empirical FFR recordings with derived FFRs, simulated via simple convolution (e.g., Goldstein and Kiang, 1958; Janssen et al., 1991; Bidelman, 2015; Carter and Bidelman, 2023). The model presumes the sustained following response is generated by a series of overlapping onset responses such that the FFR is an iterated ABR/PAM (see Fig. 1 of ref. Bidelman, 2015). We simulated FFRs by convolving each listener’s PAM-contaminated ABR (see Fig. 1) with a periodic click train of 100 Hz spacing (i.e., the FFR stimulus F0). This process generated a new ABR/PAM signature at each stimulus pulse which when strung across time, yielded a complex waveform closely mirroring the actual FFR. We then assessed correspondence between model-predicted and true FFR recordings via cross-correlation (20 ms lag search window) (Galbraith et al., 2000; Bidelman, 2015).

### Statistical analysis

We analyzed the data using a 2x2 mixed-model ANOVA in *R* (R Core Team, 2020) and the *lme4* package (Bates et al., 2015). Fixed effects were eye gaze position (5 levels) and channel montage (2 levels). Subjects served as a random effect. We used Satterthwaite’s method to compute degrees of freedom. The data were SQRT-transformed to improve normality and homogeneity of variance assumptions necessary for parametric analyses. Effect sizes are reported as 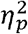.

We conducted regression analyses to assess whether a linear combination of listeners’ neuro-behavioral measures (i.e., years of music training, PAM amplitudes) predicted the strength of their FFR [e.g., FFR_rms_∼ music*PAM]. We then used leverage plots (Sall, 1990), partial correlations to assess the relative strength of music vs. artifact regressors in driving putative musicianship-FFR relations (e.g., Musacchia et al., 2007; Wong et al., 2007; Skoe and Kraus, 2012; Mankel and Bidelman, 2018). Mediation analysis was used to evaluate relationships between neural and behavioral measures. We used an efficient, bootstrapping implementation of the Sobel statistic (Sobel, 1982; Preacher and Hayes, 2004) (N = 1000 resamples) to determine whether PAM artifact fully or partially mediates the strength of listeners’ FFR. The Sobel test contrasts the strength of regressions between a pairwise vs. triplet (mediation) model (i.e., *X*_→_*Y* vs. *X*_→_*M*_→_*Y)*. Mediator *M* is said to mediate the relation between the *X*_→_*Y* if (i) *X* first predicts *Y* on its own, (ii) *X* predicts *M*, and (iii) the functional relation between *X*_→_*Y* is rendered insignificant after controlling for *M* (Preacher and Hayes, 2004).

## Results

### Postauricular muscle (PAM) artifact is present but variable across ears

We first confirmed PAM could be successfully elicited in individual ears and experimentally manipulated by directing listeners’ eye gaze towards their stimulated ear (**Fig. 1b**). **Figure 1c** shows grand average click-ABRs recorded using montages with ipsilateral mastoid vs. midline (C7) electrode reference. In both traces, eye gaze was directed toward the right (ipsilateral) ear, which was expected to elicit maximal PAM contraction (O’Beirne and Patuzzi, 1999; Patuzzi and O’Beirne, 1999). Positive deflections in the first 6 ms form the canonical waves of the auditory brainstem response (I = 1.5 ms, III=3.5 ms, V=5.5 ms) reflecting serial activation of the auditory nerve and major nuclei along the ascending auditory pathway. Following the ABR, the myogenic PAM artifact was evident between 12.5-15 ms and was especially prominent in mastoid-referenced recordings (i.e., Fpz – M2). PAM artifact was not evident at C7 where the reference electrode is distanced from the muscle. Consequently, PAM artifact was easily isolated from the neurogenic ABR by simple waveform subtraction of mastoid vs. neck recordings (**Fig. 1d**). Although PAM artifact was present in all ears to some degree, the response was highly variable, ranging from 0.18-9.7 µV (peak-to-peak amplitude; µ±σ: 2.38 ± 2.36 µV) across individuals.

### PAM contraction confounds the auditory FFR but declines with stimulus frequency and electrode montage

Having established PAM artifact was easily recordable and could be manipulated experimentally via eye gaze, we then tested whether FFR strength might also systematically vary with eye gaze direction and thus expose a myogenic source of the neural response. Confirming our intuition, FFRs to complex tones varied parametrically in amplitude when listeners were cued to direct their gaze across a ±70^0^ range (**Fig. 2a**). Hard rotation of the eyes to either the left or right ear enhanced FFRs by 3-4x, particularly between 100-200 Hz where the PAM resides within the EEG spectrum (O’Beirne and Patuzzi, 1999). FFR amplitude depended strongly on both eye gaze and recording location [position x channel:

**Figure 2:**
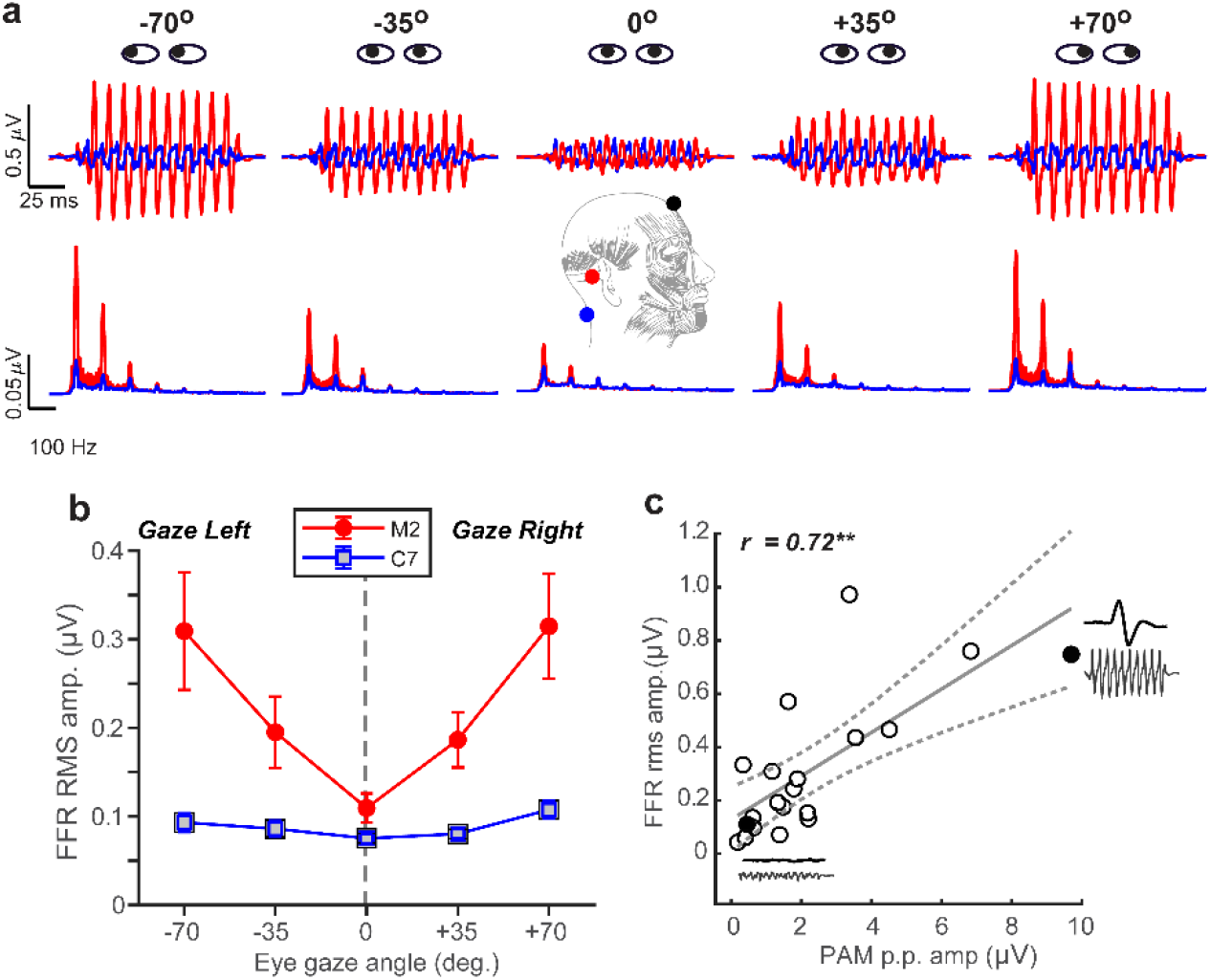
FFR strength systematically scales with the degree of PAM activation. **(a)** FFR time waveforms (*top*) and spectra (*bottom*) as a function of eye gaze spanning from left to right ear. Note the strong fundamental and integer-related harmonic frequencies of the stimulus F0=100 Hz (200 Hz condition not shown). (**b**) FFR amplitude systematically increases when gaze is directed away from the midline (in either ipsi or contra direction) for mastoid-referenced recordings. FFR amplitude is invariant to eye gaze in recordings referenced to C7, suggesting a midline montage (i.e., Fpz-C7) eradicates PAM artifact. (**c**) Correlation between PAM peak-to-peak (Fig. 1a) and FFR rms amplitudes recorded with hard right eye gaze (position #5). Solid points reflect two extreme subjects who had the largest and smallest PAM reflex (their raw waveforms are shown as insets). Dotted lines = 95% CI; errorbars= ± 1 s.e.m.; \*\**P* < 0.01

*F*_*4,152*.*16*_ = 3.72, *P* = 0.0064; 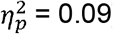]. Comparisons between the C7 and mastoid channel showed a ∼180^0^ phase shift, which is common for FFRs recorded from brainstem (highlighted in the C7 reference electrode) vs. auditory nerve (highlighted in the M2 reference electrode) (see Fig 10 in Bidelman, 2018b). For the mastoid channel, FFR strength decreased for gaze directed toward the midline (0^0^) and increased with gaze toward either ear (quadratic contrast: *t*_152_=6.82, *P*<0.0001). A nearly identical pattern was observed for the 200 Hz stimulus (data not shown; position x channel: *F*_*1, 50*.*4*_ = 8.52, *P* = 0.0052; 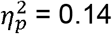). Moreover, a direct comparison of frequency effects revealed stronger FFRs in the 100 Hz vs. 200 Hz condition regardless of channel [main effect of frequency: *F*_*1, 117*.*99*_ = 4.66, *P* = 0.033; 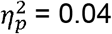], indicating PAM-related changes in FFR declined with increasing stimulus frequency. Myogenic-induced enhancements were not apparent in electrodes positioned far from the mastoid and FFRs were invariant to eye gaze when recorded from the Fpz-C7 channel (linear and quadratic contrasts: *P >* 0.17) (**Fig. 2b**).

We also found PAM peak-to-peak amplitude was highly correlated with FFR root-mean-squared (rms) amplitude [Pearson’s-*r* = 0.72, *P* = 0.0005] (**Fig. 2c**). This suggests listeners’ FFR directly scaled with the size of their muscle artifact. Collectively, these findings reveal FFR strength depends critically on eye gaze position and moreover, that PAM activity overlays true neural responses, potentially masquerading as the neurophonic FFR commonly described as having brainstem origin (Chandrasekaran and Kraus, 2010; Bidelman, 2018b; Coffey et al., 2019).

### Sustained FFRs are explained by serial PAM contractions: Model simulations

Single-unit motor neurons underlying the PAM reflex can be driven at rates up to 100-200 Hz (Kiang et al., 1963; Jacobson et al., 1964). This raises the possibility that the FFR, and what appears to be a sustained brain potential, might at least partially reflect a series of overlapping transient PAM artifacts evoked by the individual pitch pulses of low-frequency, periodic sounds. To test this possibility, we simulated FFRs via simple convolution of listeners’ isolated PAM response (i.e., Fig. 1) with a 100 Hz impulse train. This train had identical periodicity as the complex tones used in our FFR experiment. This model regenerates the PAM at each pitch period of the F0, such that responses temporally overlap and produce a quasi-steady state auditory response (Galambos et al., 1981). Repeating the PAM wavelet across time (**Fig. 3a**) yielded a sustained waveform that was strikingly similar to the empirical FFR recordings (**Fig. 3b**). Actual and PAM-derived FFRs were highly correlated (*r*_*xcorr*_ = 0.71 ± 0.21; *inset*), suggesting upwards of 50.4% of the variance (*R*^*2*^) in the FFR could be explained by a non-neuronal source.

**Figure 3:**
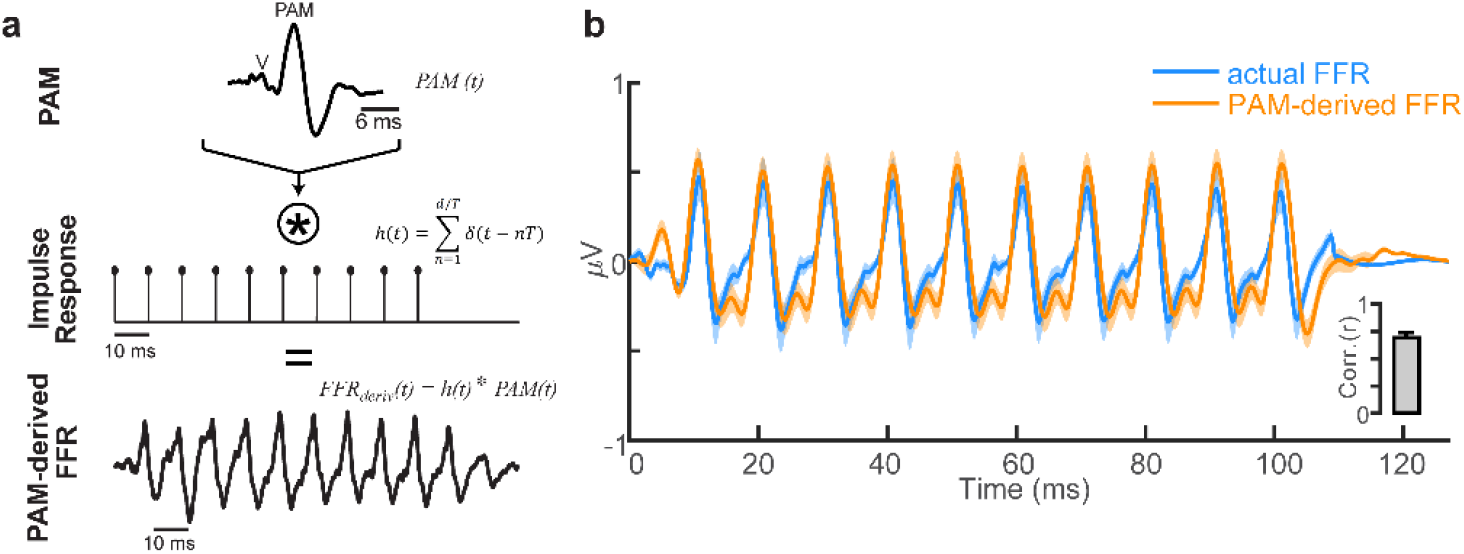
FFR waveforms can be explained as phase-locked PAM artifact. **(a)** Convolution model for generating the FFR based on repetition of the transient PAM response (a muscle artifact) (for details, see Bidelman, 2015). FFRs generated by a complex tone with period *T* are conceived as a convolution of the PAM artifact (*PAM(t)*, top row) with the individual pitch periods of the tone, modeled as a periodic pulse train (*h(t*),middle row). The resulting derived FFR (*FFR*_*deriv*_*(t)*, bottom row) is generated by a series of overlapping transient PAMs at the periodicity of the impulse train *T* = 10 ms (i.e., F0 = 100 Hz). **(b)** Comparison of empirical FFR recordings (F0=100 Hz complex tone stimulus) and simulated FFRs, derived via the convolution of PAM attract with the stimulus F0 periodicity. FFR traces reflect responses recorded with eye gaze directed at position #5 (see Fig. 1) and contain significant PAM. A series of temporally overlapping PAM wavelets repeated at the F0 of a tone accounts for ∼50% of the variance in actual FFRs (*inset*). shading/errorbars = ±1 s.e.m.

### PAM artifact mediates music-related plasticity in FFRs

Having established that FFRs are partly confounded by muscular influences, we next asked whether putative enhancements reported in the literature (e.g., due to musical training, bilingualism (Krishnan et al., 2005; Musacchia et al., 2007; Wong et al., 2007; Krizman et al., 2012; Skoe and Kraus, 2012; Mankel and Bidelman, 2018)) might be explained not in terms of auditory plasticity but rather, differences in tonic PAM contraction among listeners. To this end, we conducted multiple regression analyses to assess whether a linear combination of listeners’ neuro-behavioral measures (i.e., years of music training, PAM amplitudes) predicted the strength of their FFR [e.g., FFR_rms_∼ music*PAM]. **Figure 4** shows leverage plots (Sall, 1990) illustrating the relative effects of music training and PAM artifact amplitudes on FFR strength. Variance inflation factors (VIFs) were < 2 for both predictor variables, indicating negligible multicollinearity in the data (Lüdecke, 2020). We found that music training strongly predicted FFR strength [*r* = 0.34, *P* = 0.00076] (**Fig. 4a**), consistent with prior studies suggesting musicianship enhances the brain’s early neural encoding of complex sounds (cf. Musacchia et al., 2007; Wong et al., 2007; Skoe and Kraus, 2012; Mankel and Bidelman, 2018). However, even after partialing out musical training, the strength of listeners’ PAM remained strongly correlated with FFR amplitude [*r* = 0.49, *P* < 0.0001] (**Fig. 4b**). Similarly, musicianship was independently correlated with enhanced FFR amplitudes after controlling PAM contributions [*r* = 0.21, *P* = 0.037]. Indeed, a regression model that included the interaction between music training and PAM amplitudes was a significant improvement in describing FFR variance over one containing music alone [*F*_*1,92*_ = 29.01, *P* < 0.001]. Taken together, these data confirm that the strength of the FFR is driven by an interaction between neuroplasticity (i.e., related to listening experience) and artifactual origins.

**Figure 4:**
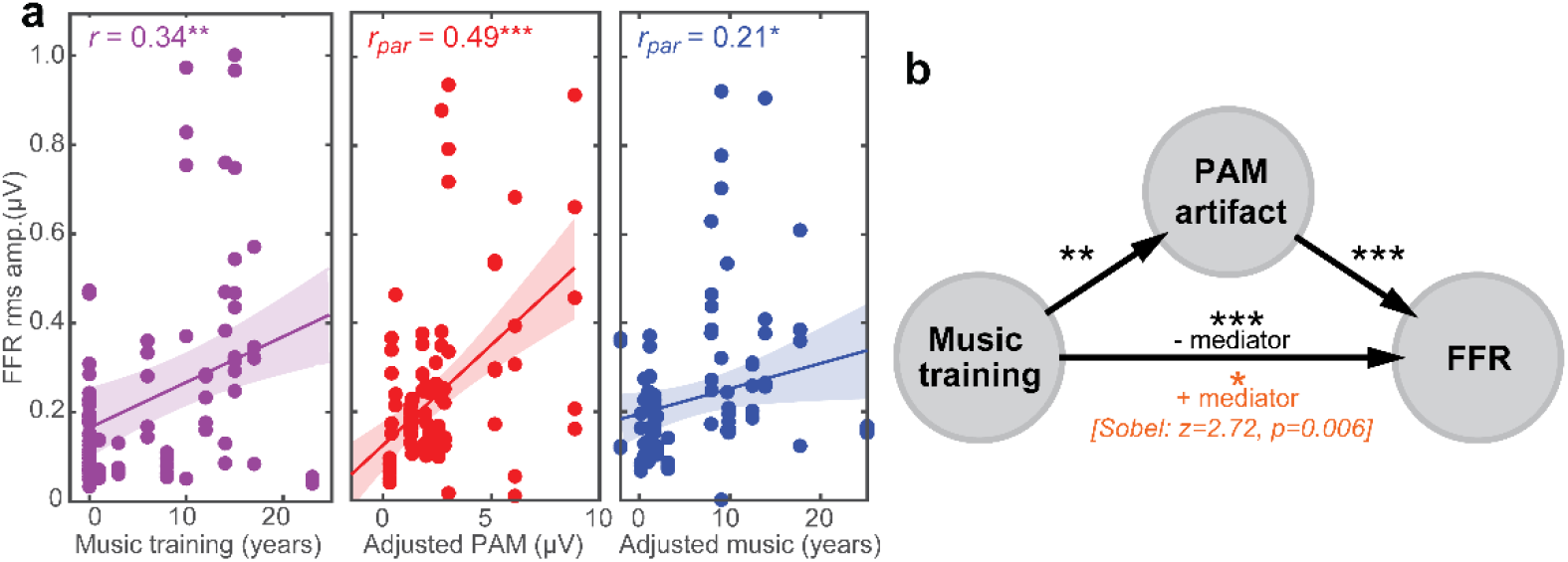
PAM artifact can masquerade as experience-dependent plasticity in the FFR. (**a**) (*left*) FFR is stronger in listeners with more musical training, suggestive of the experience-dependent plasticity reported in trained musicians (cf. Musacchia et al., 2007; Wong et al., 2007; Skoe and Kraus, 2012; Mankel and Bidelman, 2018). Both PAM (*middle*) and musical training (*right*) independently correlate with FFR strength after partialing out the other variable. Correlation data are aggregated from the 5 eye gaze positions. (**b**) Mediation analysis. The putative FFR-music relation is severely reduced after accounting for PAM [Sobel test: *z*=2.72, *P*=0.006] suggesting uncontrolled muscle artifact partially mediates the relation between musical training and FFR. Shading = 95% CI; \**P*<0.05, \*\**P*<0.01, \*\*\**P*<0.001.

To further evaluate the confounding link between PAM-related muscle noise and putative music-induced FFR plasticity (e.g., Musacchia et al., 2007; Wong et al., 2007; Skoe and Kraus, 2012; Mankel and Bidelman, 2018), we conducted mediation analyses (Sobel, 1982; Preacher and Hayes, 2004) to determine the degree to which PAM artifact mediated FFR strength (**Fig. 4b**). Mediation contrasts the strength of regressions between a pairwise vs. triplet (mediation) model (i.e., *X*_→_*Y* vs. *X*_→_*M*_→_*Y)*. Mediator *M* is said to mediate the relation between the *X*_→_*Y* if (i) *X* first predicts *Y* on its own, (ii) *X* predicts *M*, and (iii) the functional relation between *X*_→_*Y* is rendered insignificant after controlling for *M* (Preacher and Hayes, 2004). We found PAM satisfied the first two criteria for a mediating variable (**Fig. 4b**). However, accounting for the strength of listeners’ PAM artifact severely reduced—but did not render insignificant—relations between music training and FFR strength (Sobel test: z=2.72, *P*=0.006) (Sobel, 1982). These results suggest PAM artifact is a partial (but not fully) mediating variable describing music-related enhancements observed in the FFR.

### FFR timing is independent of PAM

The analyses thus far revealed links been amplitude properties of the FFR and PAM reflex. However, plasticity in the FFR has also been described in the latency of the response to speech and musical sounds, with faster neural timing following long-term musicianship (Parbery-Clark et al., 2012; Bidelman et al., 2014; Mankel and Bidelman, 2018) and short-term auditory training (Anderson et al., 2013). To explore the possibility that variation in FFR timing might also be due to undocumented PAM artifact, we examined correlations between the onset latency of the FFR [measure in the 6-12 ms search window (Mankel and Bidelman, 2018)] and PAM artifact [8-18 ms; see Fig. 1d]. However, neither latency [*r* = -0.10, *P* = 0.69] nor amplitude [*r* = 0.14, *P* = 0.57] of the PAM predicted FFR timing to the 100 Hz stimuli. Similar null correspondence was observed at 200 Hz [all *Ps >* 0.09]. Thus, in stark contrast to FFR amplitude measures, fine timing precision of the FFR appears less susceptible to artifactual influence.

## Discussion

Our findings expose a strong influence of myogenic activity on the auditory FFR. We demonstrate uncontrolled eye gaze systematically alters the apparent strength of FFRs with tandem PAM contraction. More critically, we show this muscular noise can easily masquerade as neural enhancements, accounting for large variability in the response previously ascribed to auditory-sensory plasticity. Importantly, these data do not negate the possibility that certain human experiences (e.g., music training, language expertise, learning) can confer experience-dependent plasticity that manifests in spectrotemporal changes in the FFR (Krishnan et al., 2005; Kraus et al., 2014; Mankel and Bidelman, 2018). Rather, we argue that undue muscle confounds external to the brain may play a larger role in generating the FFR and its use as a biomarker of plasticity than conventionally thought.

While typically considered an artifactual nuisance to auditory EEG and evoked potential recordings, the PAM and related myogenic potentials do find several important neuro-otological applications including hearing threshold (Yoshie and Okudaira, 1969) and vestibular assessment (Rosengren et al., 2019). Yet, in the context of FFRs whose purpose is to evaluate central auditory neural processing and the brain’s sensory representation of complex sounds, we show concomitant PAM muscle activation easily confounds these neurogenic responses, spuriously amplifying the FFR by 3-4x fold. Our findings are reminiscent of other microflexes shown to confound oscillatory EEG responses (e.g., microsaddes Yuval-Greenberg et al., 2008) and suggest uncontrolled eye movements can systematically alter sustained auditory FFR potentials via PAM muscle engagement.

We found PAM-related artifact was largest for hard rotation of the eyes toward or way from the ipsilateral ear of sound presentation but parametrically scaled in direct portion with listeners’ lateral eye gaze. While it is important to note that PAM was evoked up to a rather extreme eye gaze rotation (representing a “worst case scenario”), we also find the artifact scales in direct proportion away from midline and persists at gaze angles typical of normal eye movements. This point is particularly salient in light of speech-FFR studies that use passive listening paradigms where listeners are allowed to watch muted subtitled movies (Wong et al., 2007; Coffey et al., 2016; Mankel and Bidelman, 2018; Price and Bidelman, 2021). Videos are thought to provide added control during electrophysiological procedures, inducing a calm yet wakeful state that yields cleaner electrophysiological recordings (Wong et al., 2007; Coffey et al., 2016). However, subtended gaze angles when viewing a monitor at typical seated distance can exceed ∼30-35 degrees (Feng and Spence, 2008). Our data clearly show the FFR is nearly double its nominal size at these typical viewing angles (Fig. 2b). Consequently, while seemingly an innocuous control task, undue eye movements during video watching could easily confound the FFR by artificially inflating response amplitude. Along these lines, there is evidence that musicians show reduced fixation dwell times and more saccades during viewing tasks (Perra et al., 2022). Such increased eye movement could explain the larger PAM activation (and artifactual FFR) we find in musically trained individuals (Fig. 4). Similar augments could extend to bilinguals, who also show increased visual search (Ratiu et al., 2017) and more robust speech FFRs (Krishnan et al., 2005; Krizman et al., 2012).

While we acknowledge our gaze-induced manipulation of PAM could represent an overestimate of its activation, we note that PAM can be elicited down to hearing threshold (Purdy et al., 2005). Moreover, the strong relation between PAM and FFR amplitudes we find would remain unaffected since changes in scale affect neither the magnitude nor sign of a correlation. Thus, the fact that 50%, let alone *any* portion of variance in the FFR is explained by a myogenic artifact is a cause for concern for FFR recordings.

While there is overwhelming evidence in humans and animals to suggest PAM artifact is generated from the bipolar activation of muscle fibers behind the auricle of the ear (Kiang et al., 1963; O’Beirne and Patuzzi, 1999; Patuzzi and O’Beirne, 1999), it is conceivable that other non-auditory mechanisms contribute to FFR contamination. For example, it is well known that FFRs require delicate recording strategies to prevent inadvertent pickup of stimulus headphone artifact that can easily swamp the biological response (Price and Bidelman, 2021). Electromagnetic shielding (used here) prevents this possibility (Price and Bidelman, 2021) and can be further ruled out in the present data by the fact that FFRs showed gaze- and electrode-dependent changes that would not occur if waveforms were mere stimulus bleed (Fig. 2b). However, bogus FFR amplification could also result from other biological sources. Indeed, bone-conducted FFRs have been recorded in profoundly deaf listeners suggesting non-auditory pathways of response generation (Ribarić et al., 1984). These might include direct or indirect contributions from the vestibular system [e.g., evoked myogenic potentials (Ribarić et al., 1984; Prevec and Ribarić-Jankes, 1996; Lawlor et al., 2022)] and/or somatosensory contributions near the electrode-skin interface which can persist up to rates of 287 (Lawlor et al., 2022) and 130 Hz (Prevec and Ribarić-Jankes, 1996), respectively. Regardless of the underlying mechanism, our data clearly demonstrate physiological noise outside the auditory system easily conflates neurogenic FFR signals.

Several studies have shown enhanced voice pitch (F0) and timbre (harmonic) encoding in musically trained listeners (Musacchia et al., 2007; Wong et al., 2007; Kraus et al., 2009; Parbery-Clark et al., 2009; Kraus and Chandrasekaran, 2010; Mankel and Bidelman, 2018), though not always consistently (e.g., see Strait et al., 2012; Bidelman and Alain, 2015). Our data confirm a robust and independent relation between musical training and the strength of listeners’ auditory FFR (Musacchia et al., 2007; Wong et al., 2007; Skoe and Kraus, 2012; Mankel and Bidelman, 2018) (Fig. 4a). However, in a departure from prior studies, we show such effects are easily confounded. PAM contraction was both larger in listeners with more extended music training and partially mediated enhancements in their FFR (Fig. 4b). At first glance, these data therefore cast doubt on whether speech-FFR enhancements associated with musical training (and perhaps other experiential factors) are solely experience-driven (cf. Munte et al., 2002; Wong et al., 2007; Herholz and Zatorre, 2012) or even entirely auditory-neurogenic in nature. Undue PAM influence might explain why some studies fail to observe FFR advantages in musicians (Strait et al., 2012; Bidelman and Alain, 2015; MacLean et al., in press). Nevertheless, we show that even after controlling for PAM, music training remains a robust predictor of FFR enhancements. This indicates musicianship independently bolsters the strength of the FFR and does so above and beyond any artifactual sources.

Surprisingly, we found musicians have stronger PAM vestigial muscle reflexes than their nonmusician peers. This finding broadly converges with notions that intense auditory experiences might fortify tonic engagement of reflexive pathways that provide feedback control to peripheral hearing processes (Brashears et al., 2003; Bidelman et al., 2017b). Still, it remains to be seen whether other long-term listening experiences that also enhance FFR similarly elevate PAM and account for neuroplastic effects observed in bilingualism and tone-language expertise (Krishnan et al., 2005; Krizman et al., 2012; Zhao and Kuhl, 2018). Regardless, the intrusive nature of PAM mandates caution for interpretating even short-term changes in the FFR, as observed in rapid perceptual learning studies (Anderson et al., 2013; Reetzke et al., 2018). More successful learners could show increased arousal while acquiring novel auditory information, leading to increased eye movements that exaggerate PAM artifact, and in turn false amplification of FFRs. FFRs are modulated by listeners’ arousal state at the single-trial level (Lai et al., 2022; Carter and Bidelman, 2023) and eye movements change during the learning process (Laamerad et al., 2020). Thus, while it awaits empirical confirmation, our data raise the possibility that rapid plasticity observed in FFRs during short-term auditory/speech learning tasks might be due to not improvements in the brain’s sensory representation of sound, *per se*, but an unmeasured artifactual source (Anderson et al., 2013; Reetzke et al., 2018).

Fortunately, our data offer a direct solution to easily thwart PAM-related confounds in the FFR and allow for untainted assessment of neuroplasticity. FFRs are frequently recorded using a differential electrode montage and non-inverting reference electrode placed on the bony mastoid process of the ipsilateral ear of stimulation (e.g., Fpz-M1/M2; see Fig. 1c, inset). We show this common practice is highly problematic, leading to extraneous pickup of the adjacent PAM muscle and overinflation of FFR that, by our estimates, has affected nearly half the published literature (see Fig. 1a and *SI Materials*). Repositioning electrodes to the earlobe (another common technique) (Chandrasekaran and Kraus, 2010; Skoe and Kraus, 2010) attenuates but does not fully eliminate PAM pickup (O’Beirne and Patuzzi, 1999). Instead, we find clean FFRs are easily recorded with electrodes placed on the upper neck (C7 vertebra), distal to the pinna and PAM muscle fibers. A midline Fpz-C7 montage has the further advantage that it (i) reduces pickup of more peripheral (cochlear) sources of the FFR and (ii) is optimally oriented with the vertical dipolar sources in the brainstem that dominate generation of the neurogenic FFR (Galbraith et al., 2000; Bidelman, 2015; 2018b). At present, it appears only 16% of studies use this optimal C7/neck montage (Fig. 1a). Secondly, we showed that FFR latency measures were largely independent of artifactual influences. Thus, timing characteristics of the FFR might provide a more veridical index of neuroplasticity (Parbery-Clark et al., 2012; Anderson et al., 2013; Bidelman et al., 2014; Mankel and Bidelman, 2018) than amplitude measures alone when tracking brain changes due to experience-dependent factors and novel sound learning (but see Mankel and Bidelman, 2018). Thirdly, controlling participants’ eye gaze straight forward (e.g., with a fixation cross) or instructing them to close their eyes could help minimize PAM contributions to the FFR. Still, we note that changes in eye gaze within only a small viewing angle could affect FFR and eye closer does not prevent occular movement altogether as the eyes are still free to move under the eyelid. Lastly, we found that PAM-related changes in FFR declined with increasing stimulus frequency. This suggests that in addition to isolating brainstem-centric sources of the response without cortical contributions (Bidelman, 2018b), the use of auditory stimuli with higher F0s (e.g., > 200 Hz) might be further advantageous in protecting against PAM confounds in the FFR.

In conclusion, our data provide new evidence that myogenic sources can artificially inflate the sound-evoked FFR and might account for certain *de novo* enhancements in the response. Provocatively, they also show that formally trained musicians have additional gains in sensory brain processing above and beyond PAM-related confounds. Our results emphasize the critical need for future studies to safeguard against ocular and myogenic confounders before claiming changes in FFR track with improvements in auditory brain function due to experience-dependent factors or neurobehavioral interventions. Of practical implication, we recommend that to measure true neurogenic FFRs without undue electromyographic contamination, investigators should (i) adopt a midline, necked-referenced recording approach, (ii) use high-frequency F0 stimuli (> 200 Hz) where PAM contributions are minimal, and (iii) focus primarily on response latency quantification to index disordered auditory processing and/or neuroplastic changes due to intervention.

## Acknowledgments

Work supported by the National Institutes of Health (NIH/NIDCD R01DC016267).

## Data availability

Data associated with this manuscript are provided in the *Supplemental Material*.

## Footnotes

1 Sohmer et al. (1977) describe two cases studies from “normal subjects with an unusually large response from the post-auricular muscle (p.656).” They observed, “the amplitude of the post-auricular responses to click stimuli and this frequency following of the post-auricular muscle were both susceptible to head position, that is to the degree of neck tension (p.660)” and concluded the PAM contributes to the FFR. Unfortunately, no quantitative analysis was undertaken so these observations remain qualitative.

2 In practice, 70^0^ rotation is difficult to achieve as human eye movements rarely exceed 50-60^0^ (Lee et al., 2019). In these extreme conditions, listeners simply panned their eye gaze left/right as far as possible.

