## Supplementary material for "Myogenic artifacts masquerade as neuroplasticity in the auditory frequency-following response (FFR)": Suppmental Data (from another version of this preprint: version 1, 2023-11-01)

### Supplemental references (N=314 FFR studies; 1973-2023)

33. Bhagavan SG, Kalaiah MK. (2020).

106. Galambos R, Hecox K. Clinical applications of the human brainstem responses to auditory stimuli (NASA-CP-143134). In: *Proceedings of the Brussels Conference* (ed^(eds Desmedt J). NASA Center for Aerospace Information (CASI) (1975).

230. Plack CJ, Fitzpatrick S, Carlyon RP, Gockel HE. A Temporal Code for Huggins Pitch? In: *The Neurophysiological Bases of Auditory Perception* (ed^(eds) (2010).
